# Bacterial cellulose spheroids as building blocks for 2D and 3D engineered living materials

**DOI:** 10.1101/2020.05.11.088138

**Authors:** Joaquin Caro-Astorga, Kenneth T. Walker, Tom Ellis

**Affiliations:** Department of Bioengineering, Imperial College London, London, SW7 2AZ, UK; Imperial College Centre for Synthetic Biology, Imperial College London, London, SW7 2AZ, UK

## Abstract

Engineered living materials (ELMs) based on bacterial cellulose (BC) offer a promising avenue for cheap-to-produce materials that can be programmed with genetically encoded functionalities. Here we explore how ELMs can be fabricated from millimetre-scale balls of cellulose occasionally produced by *Acetobacteriacea* species, which we call BC spheroids. We define a reproducible protocol to produce BC spheroids and demonstrate their potential for use as building blocks to grow ELMs in 2D and 3D shapes. These BC spheroids can be genetically functionalized and used as the method to make and grow patterned BC-based ELMs to design. We further demonstrate the use of BC spheroids for the repair and regeneration of BC materials, and measure the survival of the BC-producing bacteria in the material over time. This work forwards our understanding of BC spheroid formation and showcases their potential for creating and repairing engineered living materials.

## Introduction

Engineered living materials (ELMs) are those containing cells on or within the material that play a role in its functionalization or can produce the material itself^1,2^. Bacterial cellulose (BC) is a carbohydrate polymer produced by many bacterial species as a structural element of their biofilm and offers excellent opportunities for developing new ELMs^3^. The BC materials produced by several *Acetobacteriacea* species are of particular interest as these are quickly and cheaply made as pellicles - a large mass of thick BC - when the cells are grown in static rich media^4,5^. Bacterial cellulose inherently has attractive mechanical properties and crystallinity, has a high water-retention capacity and is ultra-pure compared to plant cellulose^6,7^. These outstanding properties of BC make it an excellent candidate for developing new materials with improved technical and environmental benefits. In the last decade, progress in understanding and producing BC has now led to its use in a broad range of applications, including products used in textiles, cosmetics, healthcare, audio-visual technology and architecture^8–11^. Most of these applications use sterile, purified BC as a bulk specialised material, however bacterial cellulose has also shown promise as an ELM^12,13^. In one recent example, incorporating *Bacillus subtilis* cells into BC-based wound dressings helped to prevent wound infections by blocking the growth of several pathogenic bacteria^14^.

Two desirable features of ELMs not routinely seen in normal materials are regeneration in response to damage, and modular design with patterned functionalities. Easy and cheap repair of damaged materials (or their automatic regeneration) is an important consideration for the sustainability of all new materials^15^. BC offers excellent opportunities in this regard, because the bacteria trapped in the grown material have the potential to regenerate it by further growth and cellulose production in the future. Just by providing nutrients, water and oxygen, the bacteria in theory can keep growing and seal gaps and tears when they arise, so long as the material has not been sterilised after growth. For patterned functionalities, this can also theoretically be achieved with BC-based materials by growing these from genetically engineered cells programmed^3^. However, another possibility to tackle this problem is to use modular ELM building blocks and pattern these physically to make larger materials. Such a ‘building block’ approach to novel materials has been taken before in nanotechnology to increase the complexity of materials and to facilitate industrial scaling of complex pieces^16^. Modular BC-based building blocks have not been explored before in an ELM context, but BC and in particular its rapid production from living cells within the material structure, offer an excellent opportunity to tackle this challenge.

Past work has shown several solutions for building BC into shapes other than the standard flat pellicles. Growing BC in hyper-hydrophobic moulds have allowed researchers to create a versatile range of three dimensional (3D) shapes with high accuracy^17,18^. However, moulds are limited in what they can achieve and typically work just for growing one material at a time. Creating patterns of functionalised BC grown from several different cell types would prove difficult with this approach, and so limits its use for creating 3D BC-based ELMs. 3D-printing of cells with semi-solid growth support materials is more promising in terms of creating 3D BC-based ELMs incorporating multiple strains in patterns^19^. However, the additive manufacturing approach relies on specialised equipment and requires washing the printed matrix to obtain the final shape. A building-block approach where modular units grow and self-connect into 3D shapes would be less cumbersome.

Here, we introduce the concept of using a building block approach to make 3D BC-based ELMs. This is achieved by growing and utilising BC ‘spheroids’ - millimetre-scale spheres of actively growing BC that have been observed in prior work. Growth of BC spheroids has remained poorly-understood and is typically characterised as strain-dependent or inconsistently produced^20^. Here, we now define a reproducible method to produce BC spheroids from the bacterial strain *K. rhaeticus* iGEM. We use these BC spheroids as building blocks to build complex 3D shapes and to create mosaic patterns made of spheroids containing genetically functionalized bacteria that impart fluorescence. We further demonstrate that spheroid building blocks can be used to regenerate damaged BC materials, growing and interweaving with an existing BC piece.

## Methods

### Strains, culture conditions and BC pellicle growth

The *Acetobacteriacea* strain used in this work was *Komagataeibacter rhaeticus* iGEM. The wild-type version of this strain was tagged with green and red fluorescence by transformation with the plasmid KTK_124 and KTK_182, These plasmids expresses superfolder green fluorescent protein (sfGFP) and red fluorescent protein from the J23104 constitutive promoter. They were constructed by Golden Gate assembly modifying the BioBrick compatible vector pEMpty (previously pSEVA331Bb; *E. coli K. rhaeticus* expression vector, ori pBRR1 origin of replication, chloramphenicol resistant)^21^ Bacteria were transformed by electroporation using the *E. coli* protocol as described in Florea *et al*^21^.

Pre-cultures of *K. rhaeticus* were prepared by taking cells from −80°C stocks and growing in 50 ml tubes with 10 ml of Hestrin and Schramm (HS) media (peptone 5 g/L, yeast extract 5 g/L, 2.7 g/L Na_2_HPO_4_, 1.5 g/L citric acid, 2% glucose, 2% cellulase from *Trichoderma reesei* [Sigma-Aldrich]) in shaking conditions at 250 rpm/min and at 30°C for 3 days. 2 x HS media was used for spheroid production tests and was prepared by doubling the concentrations of peptone and yeast extract.

To grow pellicles, pre-cultures were centrifuged at 7000 g for 3 min, cells were then resuspended in 10 ml of HS without cellulase. This process was then repeated. The suspension was diluted 1 in 100 to grow pellicles, growing in shallow containers with 200 ml of HS without cellulase, supplemented with 1% ethanol and incubated at 30°C for several days. To grow cellulose spheroids, the suspension was diluted 1 in 1000 in 3 ml of HS without ethanol, in 15 ml tubes, in shaking conditions at 250 rpm/min and 30°C for 3 days.

### Bacterial cellulose synthesis time-lapse

A frozen glycerol stock of *K. rhaeticus* was used to inoculate 5 mL of HS media containing 2% glucose. The culture was incubated static for 7 days at 30°C until a pellicle formed. To prepare the microfluidic plate, 50 μL of culture from underneath the pellicle was removed and placed into the inlet well of a B04A CellASIC ONIX plate (Merck). The plate was placed onto a fluorescence microscope (Nikon Eclipse Ti inverted microscope) within a chamber heated to 30°C, and cells were fed with a continuous flow at 1 Psi of HS media containing 2% glucose and 0.001% Fluorescence Brighter 28 (Sigma-Aldrich). After 24 hours, growing cells were identified and a time-lapse was started using the bright field and DAPI channels, and images were taken every 2 minutes for 2 hours.

### 3D structures and fluorescence

Spheroids from day 3 cultures of both wild-type and sfGFP-tagged strains were collected by filtering the culture with filter paper in sterile conditions. The spheroids were seeded in the desired shape with the help of a pipette tip and then incubated for 4 days at 30°C. To create patterns, spheroids were taken one-by-one using sterile pipette tips and placed together in the desired positions.

Pellicles repaired with fluorescent spheroids and spheroid patterns made with fluorescently tagged cells were imaged with an Amersham Typhoon Scanner, using 10 μm resolution. A far-blue light gel transilluminator with amber filter was used to image spheroids produced by sfGFP-tagged cells.

### Pellicle repair

A 0.5 cm hole puncture tool (Jenley hollow leather punch) was used to create holes in bacterial cellulose pellicles. For repair using spheroids, spheres from day 3 cultures were filtered with filter paper in sterile conditions to separate them from the liquid culture. The spheroids were placed in the puncture holes and 50 μl of HS supplemented with 2% glucose was dropped over the spheroids. 2 ml of HS was then added into the surrounding container of the pellicle to maintain it in a hydrated state. The pellicles were incubated in static conditions for 10 days at 30°C before imaging. In unsuccessful attempts of pellicle repair the following were used i) fragments of biofilm found adhered to the wall of a flask after 4 days in shaking conditions; ii) floating clumps formed in shaking conditions from a initial culture set with high cell density (OD_600_ ~0.5); iii) cellulose aggregates present in the culture medium under the pellicle; iv) a pellet of cells grown in shaking conditions with cellulase, centrifuged, washed with HS and centrifuged again; v) cells from (iv) embedded in a 0.3% HS agar matrix at 40°C and immediately placed in the pellicle before it solidifies; vi) a cellulose patch of slightly bigger dimensions than the hole produced in the pellicle, used to force the edges of the patch and the hole to be in close contact.

### Cell survival in pellicles

A homogeneous suspension to grow pellicles was prepared as described above. Pellicles were grown in 96 squared deep well plates at 30°C. After 7 days, pellicles were stored in vacuum sealed plastic bags at 4°C and 23°C and also in 2 ml tubes at 23°C. Samples in triplicate were collected at each time point to assess survival. Pellicles were placed in 2 ml tubes, suspended in HS diluted 1 in 10 with 5% cellulase and incubated at 37°C for 3 h in shaking conditions to degrade the cellulose. Serial dilutions of each suspension were made, and these dilutions plated in four replicates on HS agar supplemented with 2% of glucose. After 7 days of incubation at 30°C, colonies were counted from the agar plates and colony forming units (CFUs) per cm^2^ pellicle area was calculated.

## Results

### Bacterial cellulose spheroids

Bacterial cellulose (BC) spheroids have been reported in several previous works^20,22,23^, but how they form and why has not been fully elucidated. It has been hypothesized that spheroid formation is produced by the adhesion of bacteria to air bubbles produced in shaking media, with cellulose then grown at the bubble air-liquid interphase to form a spheroid shape^24^. Past experiments in our lab growing *K. rhaeticus* in shaking conditions occasionally produced spheroids. As with other BC-producing bacteria, we originally assumed that spheroid formation by *K. rhaeticus* occurred randomly, either in response to stochastic processes during shaking growth or due to mutation or another form of uncontrolled variation in cell behaviour that triggers their spontaneous production.

Here we set out to examine whether spheroid production by *K. rhaeticus* was indeed a random event, or one that could be reproducibly triggered. To do this, we grew *K. rhaeticus* cells with shaking at 30°C and tried combinations of more than 20 different growth variable (Supplementary Table 1). During and after growth, we visually assessed the cultures for the presence of BC spheroids and used this information to determine the key factors involved in spheroid formation and which combination of growth variables leads to reproducible growth of BC spheroids.

Our results indicated that the main determining factor for BC spheroid formation is the initial optical density of the culture, with more ideal spheroids seen when cultures begin at low optical densities (OD_600_ = 0.001-0.0001) where bacteria are more likely to begin isolated from one another (Supplementary Figure 1A). The second most important factor seen in our experiments was the culture container. BC spheroids were only ever seen forming after shaking growth in 14 ml and 15 ml plastic culture tubes, and never grew in any of our attempts with 50 ml tubes or with larger bottles and flasks of different sizes, regardless of the different culture volumes tested. The third factor revealed in our experiments was the culture media. The addition of 1% ethanol is known to drastically increase the BC yield when growing static pellicles^25^. Unexpectedly, however, we found in multiple trials with different seeding dilutions and containers that ethanol inhibits spheroid formation. The use of 2 x HS instead of normal HS media gave a higher yield in the number of spheroids per tube, but the spheroids were smaller in size and less uniform (Supplementary Figure 1B).

With these three factors determined, it became possible for us to produce a protocol for reliable spheroids production (Figure 1A) in 14 ml culture tubes, yielding spheroids typically 0.2 to 1 mm in diameter (Figure 1B). We also tested if our protocol for spheroids cultivation would also work for genetically modified strains of *K. rhaeticus* containing plasmids expressing transgenes. We observed reliable growth of green fluorescent spheroids from *K. rhaeticus* expressing sfGFP and red fluorescent protein (RFP) expressed from a plasmid (Figure 1C).

**Figure 1.**
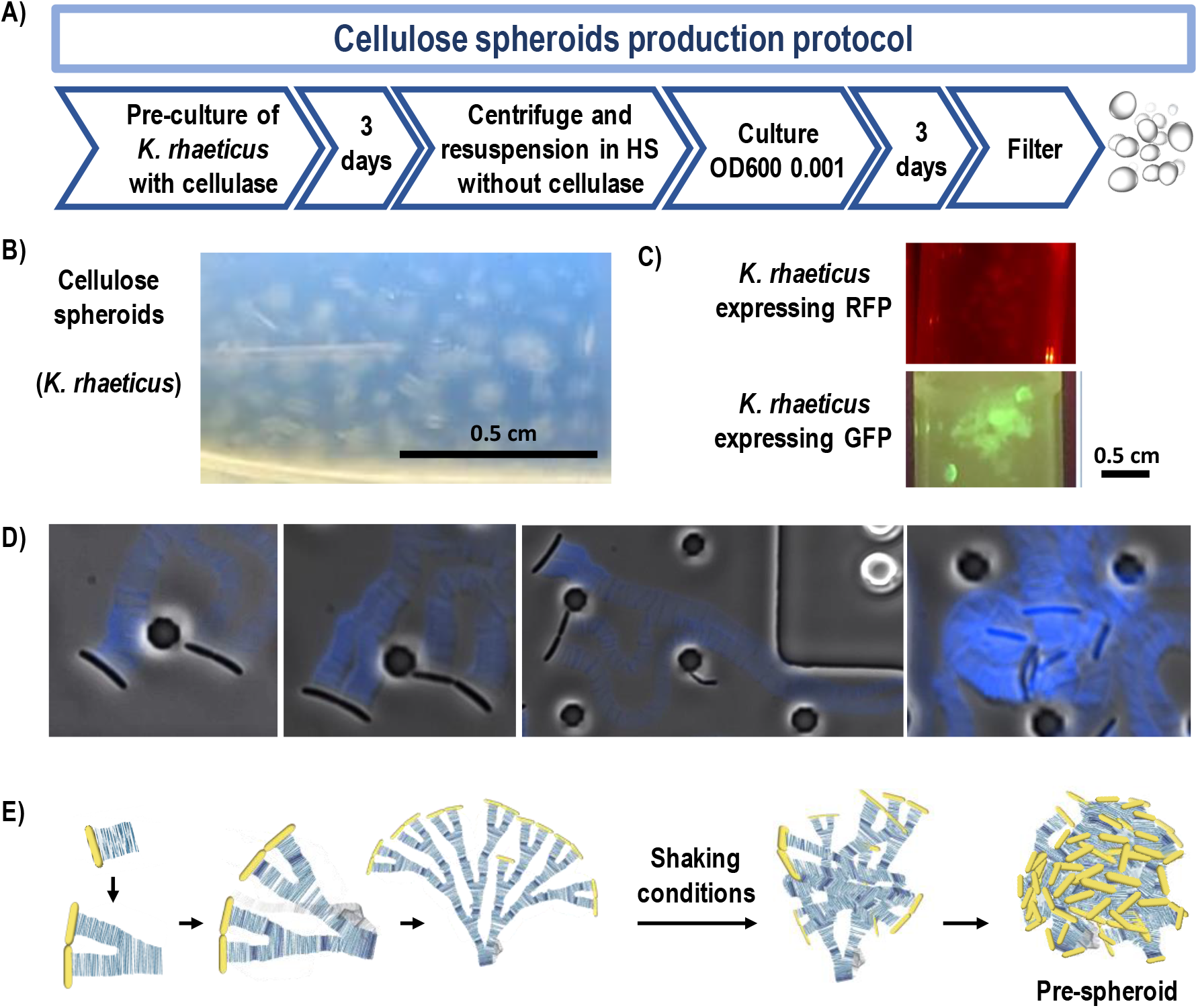
Bacterial cellulose (BC) spheroids. A) Schematic of the protocol to produce BC spheroids from *K. rhaeticus* in shaken cultures. B) Example image of BC spheroids produced by wild type *K. rhaeticus* cells. C) Example image of BC spheroids produced by sfGFP (left) or RFP-tagged (right) *K. rhaeticus* cells. D) Microscopy images of a microfluidic growth time-lapse of *K. rhaeticus* cells growing at low density with calcofluor blue staining of cellulose. E) Schematic showing growth progression of cellulose in static and shaking conditions, hypothesised to lead BC spheroid formation.

In order to observe how BC is produced by our bacteria, we performed microfluidic time-lapse culture experiments with calcofluor added to stain nascent cellulose production (Supplementary Video 1). We observed band-like growth of BC chains coming from one side of the bacteria longitudinal axis, producing branches of cellulose as the cells divide (Figure 1C). In static culture, this event is responsible for the formation of layers of cellulose growing into pellicles. However, when perturbed by shaking, the branches collapse on themselves, entangling the BC bands while the chains continue growing and cells dividing. Although conditions in the microfluidic chamber do not match those used for spheroid production in our protocol, we reason that the same processes lead to the formation of spheroids when cultures are seeded at very low density (Figure 1D).

### Construction in 2D and 3D with BC spheroids

Given their mechanism of growth, we reasoned that spheroids would continue cellulose production at their surfaces and thus when two spheroids interact for enough time they will grow together and fuse. This property of our spheroids would allow them to act as millimetre-scale BC-based building blocks that could then be used to produce 2D and 3D shapes (Figure 2A). To demonstrate this application, we designed a simple 3D shape (a podium) and a more complex 3D shape (a serrated ring). We then manually placed spheroids in these arrangements on sterile cotton pads, with the help of a pipette tip. After 4 days of incubation at 30°C, the spheroids had visibly grown and fused together to create a continuous BC-based shape roughly matching the seeded design (Figure 2B).

**Figure 2.**
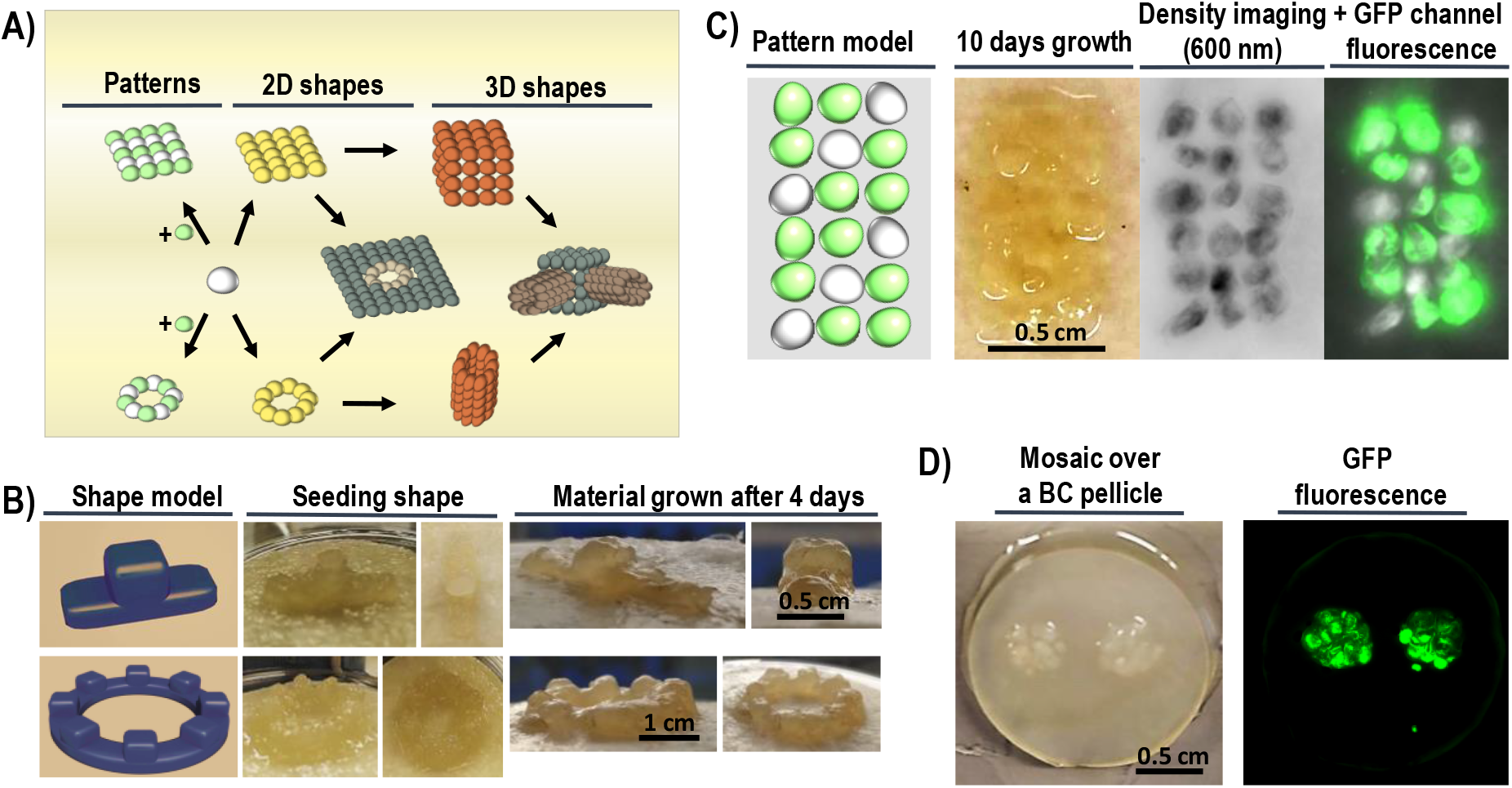
BC spheroids as building blocks. A) Schematic of potential building structures that can be created from spherical building blocks. B) Growth of simple and complex 3D shapes constructed using BC spheroids. Model (left), spheroid seeding (middle) and result after 4 days of growth (right). C) Patterns of functionalized BC spheroids. Model (left), pellicle after 10 days growth (middle left), spheres imaged using 600 nm excitation laser (middle right) and merged 600 nm image and GFP channel. E) Mosaic of sfGFP-tagged spheroids grown for 4 days at 30°C on an existing bacterial cellulose pellicle. Pictures of the material after 10 days of growth in the visible channel (left) and green fluorescence channel (right).

The use of cellulose spheroids as building blocks opens a new opportunity to create ELMs with 2D and 3D patterns and multifunctional properties. As *K. rhaeticus* is a non-motile bacteria, the patterns created from spheroid seeding will not get blurred and should remain conserved. To demonstrate, we designed and created layer of BC seeded onto filter paper with both normal BC spheroids and BC spheroids made by GFP-tagged *K. rhaeticus*. We placed the fluorescent spheroids in three lines close to each other, setting a pattern to make diagonal lines of non-fluorescent spheroids within the pattern. After 10 days of incubation at 30°C, we obtained a fused pellicle conserving the fluorescent pattern of seeding, demonstrating the possibility of easily creating ELMs functionalized at the millimetre scale - the diameter of a single spheroid (Figure 2C).

We observed in these experiments that growth after seeding also led to the spheroid structure being adhered to the sterile filter paper support. As paper is itself predominantly cellulose, we wondered how the spheroids would behave if they were set to grow on a layer of bacterial cellulose. To examine this, we seeded fluorescent spheroids on a normal *K. rhaeticus* pellicle produced from static growth. After 4 days of incubation at 30°C, the spheroids were completely fused to the base pellicle, revealing that a BC layer provides an excellent frame within which to set spheroids in patterns. Fluorescence was localized exactly in the area of seeding (Figure 2D).

### Bacterial cellulose regeneration by BC spheroids

Given the strong interest in using BC as a basis for ELMs, there is a need to identify methods to repair or regenerate a BC-based material when and if they are damaged. The efficient fusing of spheroids into a pellicle as seen in pattern formation experiments (Figure 2D) suggests that spheroids could provide a method for pellicle damage repair (Figure 3A). To investigate this, we established a repair assay of BC pellicles BC using a hole punch to damage the material. We first assessed whether just the addition of HS media and further incubation for 7 days in static aerated conditions would result in regrowth of BC in the hole wound of a punctured fresh BC pellicle. Although a thin BC layer did grow over the hole, the new thin pellicle layer was poorly adhered to the original one below it (Supplementary Figure 2A). We considered that the poor adherence was a result of adding too much liquid growth media, inducing the formation of a new BC layer only at the air liquid interface and not well-adhered to the original pellicle. Thus, we decided to just add a few drops of new HS media in the holes with and without also adding the circles of cellulose produced by the hole puncture (Supplementary Figure 2B, ‘Controls’). After 7 days incubation, this still failed to show stable wound repair.

**Figure 3.**
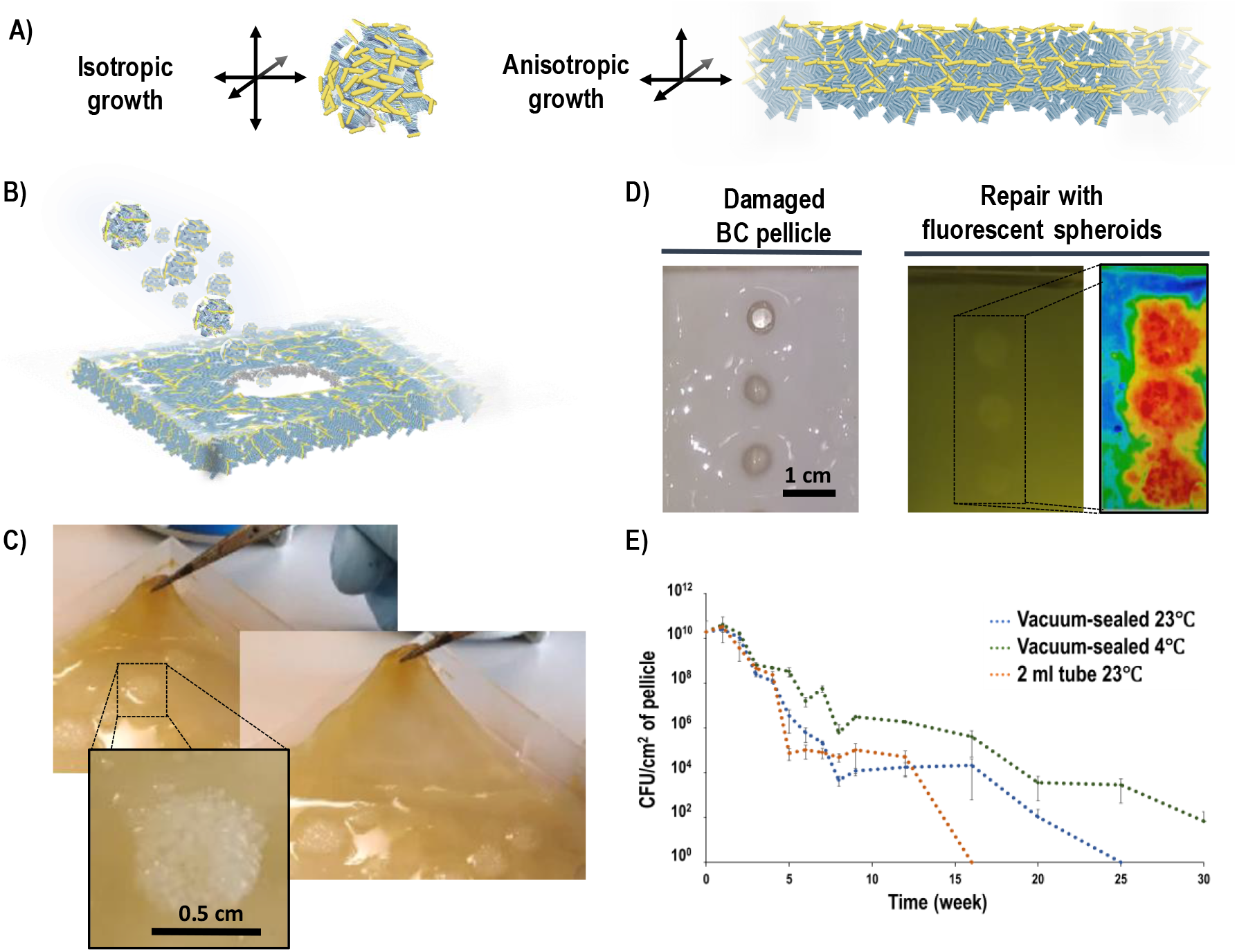
Repair of BC materials using BC spheroids. A) Illustration of the BC growth axis of spheroids compared to those seen in pellicles. B) Schematic of the regeneration strategy of repair by seeding puncture holes with spheroids. C) Result of the regeneration of a bulk BC pellicle obtained using BC spheroids. Light passing through the regenerated material layer does not suffer diffraction, indicating recovery of the bacterial cellulose macrostructure. D) Result of the regeneration of a sterilised BC pellicle (left) using spheroids of *K. rhaeticus* cells tagged with sfGFP (right) and incubated for 10 days. The heat map indicates intensity of green fluorescence, revealing the greatest intensity inside the seeded spheroids and weaker but measurable signal corresponding to the colonization of the local region of pellicle. E) Graph of *K. rhaeticus* cell survival over time from within BC pellicles stored in vacuum-sealed plastic bags or in 2 ml tubes at 4 and 23°C. Plotted values are the mean of 3 biological replicates of 4 technical replicates each.

This failure to repair may be explained by the fact that BC pellicles grow anisotropically; producing new cellulose predominantly in the horizontal axis at the top layer, building the pellicle from the bottom up by stacking cellulose layers one over the other. In contrast, spheroids grow BC isotropically, producing it in every radial direction (Figure 3A). We have observed that the incubation of two stacked pellicles does not produce regrowth and their fusion, so next we looked to see how spheroids can behave as a repair material. After placing freshly-grown spheroids into the puncture hole at high density (Figure 3B) and incubating for 6 days to allow for cellulose growth, we saw excellent repair that was not only stable but also restored the consistency and appearance of the top layer of the material (Figure 3C). We also compared this repair method to many other options for repair where growing cellulose-producing bacteria or fragments of pellicles in different forms are placed into the wound holes and allowed to grow (Methods and Supplementary Figure 2B). Repair quality was assessed by holding the pellicle edges with tweezers and pulling. Only the pellicle restored with cellulose spheroids maintained the continuity of the pellicle with high enough quality to be stable upon further manipulation (Figure 3C). We speculate that this is due the growth axis of the materials used for restoration.

Notably, the BC layer repaired with spheroids did not look different to an unpunctured pellicle once the repaired pellicle was lifted. This suggests that there is no change in the diffraction angle of the light between the original BC and newly synthetized cellulose from the spheroids, perhaps due to the spheroids producing similar cellulose that infiltrates and networks with the existing BC material (Figure 3C). To further investigate how spheroids interlink with a BC pellicle, we performed a further repair assay experiment, but now refilling the wound holes with fluorescent spheroids produced by GFP-tagged bacteria. After 10 days, the holes were repaired, and fluorescence could be observed within the hole seeded with spheroids and also at a weaker level in the surrounding material of the pellicle (Figure 3D). This reveals that bacteria spread into the local region of the damaged cellulose sheet into which they are placed.

In parallel to studying repair using BC spheroids, we also investigated how long unsterilized BC-based materials maintain the population of *K. rhaeticus* cells within them. This was assessed in order to understand whether BC spheroids could be made in bulk in advance, stored and then used for construction or repair when needed. Assessing cell survival within the spheroids proved difficult due to the small size of spheroids and irregular shapes giving variable volumes and thus cell counts. Therefore, we instead used small BC pellicles grown in 96 well microplates as an equivalent material. After growth of this BC material, the small pellicle samples were immediately stored in either vacuum-sealed bags (stored at 4°C and 23°C) or in 2 ml tubes at 23°C. Then over a period of time, the samples were removed from storage, digested with cellulase and grown on solid agar plates in order to determine cell number per sample by calculating colony forming units. This revealed that the number of viable cells in the material decreased rapidly in the first 3 weeks for all three storage methods tested (Figure 3E). However viable cells were still recovered beyond 12 weeks in all cases, and when the BC was vacuum-sealed and stored at 4°C, cells were still recoverable beyond 6 months.

## Discussion

In the research presented here, we have uncovered the basic principles of BC spheroids formation from growing *K. rhaeticus* iGEM cells and have determined a reproducible protocol for their production. Previous research on BC spheroids has suggested that their production is strain dependent with some strains capable of their production but others not. For example, the *G. xylinus* JCM 9730 strain (ATCC 700178) was shown to produce spheroids but *G. xylinus* NCIMB strain (ATCC 23769) did not^20^. Here we show that BC spheroids can be routinely made from *K. rhaeticus* iGEM and that a by having a consistent low number of cells to begin the culture is critical. It may be the case that all BC-producing bacteria can produce BC spheroids if seeded at the correct density. We note that the protocol of the aforementioned work did not measure the cell density of the bacteria inoculum before beginning culturing for BC spheroid production.

While previous work has hypothesised that attachment and growth around air bubbles triggers spheroid production^25^, our data from microfluidic time-lapse imaging offer a new insight, showing that there can be an entangling process of cells and cellulose chains during the early stages of culture growth. As uncovered here, BC spheroid formation from *K. rhaeticus* iGEM strongly depends on the bacteria concentration used to seed the culture, with low culture density being required. When the shaking culture starts at high density, we always obtain amorphous aggregates of cellulose that range from a very fibrous mass to single large piece of rounded cellulose. This tallies with previous work with *G. xylinus* JCM 9730 that showed that differences in the number and size of spheroids depended on the volume used to inoculate the culture^26^. The volume and shape of the container also affected spheroid formation, likely due to how the motion of the growth media is affected and how this changes the entangling process and air bubble formation.

Our work also demonstrated the potential for BC spheroids to be used as building blocks to create 2D and 3D shapes and create patterns that could find use as functionalized ELMs. The building block approach opens a myriad of applications, especially when considering the possibility of using BC spheroids grown from genetically reprogrammed cells that perform other tasks. Through synthetic biology, bacteria can be made to sense a wide variety of biological, chemical and physical inputs and also to signal to one another in ways analogous to electric circuitry^27–29^. Different functionalized spheroids could be used to create 2D and 3D patterns that compute environmental information, or even patterned materials that display anchor proteins to attract mammalian cells or cell differentiation signals. Such a material could for example be used to seed and grow complex mammalian tissues like skin or cartilage in defined patterns and layers.

BC spheroids also offered the best solution to BC material repair in the research shown here. Regeneration of a damaged BC pellicle and restoration of its continuity was seen in only a few days of growth with seeded spheroids. Our best results were obtained with thin pellicles. For thicker pellicles, we observed that restoration was superficial, and we speculate that this is due to the poor penetration of oxygen to the spheroids in deeper layers. To solve this problem, we now perform a double incubation, turning over the pellicle after a few days to also let the material to heal on the other surface. BC spheroids function as a successful vector to seed bacteria for BC repair and our data suggest they may be able to be stored for several months prior to use, although likely with reduced repair capacity over time.

For creating small 3D shapes, BC spheroids building blocks are limited by their millimetre size and the low precision of fabrication by hand. For making smaller or more precise BC-based ELM shapes it may be more convenient the use 3D printing methods developed for bacterial cultures, such as the FLINK method where a non-living gel matrix that harbours the bacteria is printed into the desired shape and the cells then grow and produce within this^19^. 3D printing allows the production of functionalized ELMs using different bacteria, but its accuracy is limited by the width of the extrusion printing head, which itself is limited by the high viscosity of the printed gel. A possible solution is a hybrid approach where a 3D printer is programmed to precisely dispense BC spheroids, and the size of these spheroids is reduced by harvesting them a day earlier than in the protocol given here.

## Acknowledgements

The authors wish to thank Dr Koonyang Lee, Dr Vivianne J Goosens and Mr Amritpal Singh for their help with this project at Imperial College London. We acknowledge the US Office of Naval Research Global (ONRG) and US Army CCDC DEVCOM grant W911NF-18-1-0387, and the UK Engineering and Physical Sciences Research Council (EPSRC) grant EP/N026489/1 for funding this work.

## References

1. Nguyen, P. Q., Courchesne, N.-M. D., Duraj-Thatte, A., Praveschotinunt, P., & Joshi, N. S. Engineered Living Materials: Prospects and Challenges for Using Biological Systems to Direct the Assembly of Smart Materials. Adv. Mater. 30, 1704847 (2018).

2. Chen, A. Y., Zhong, C., & Lu, T. K. Engineering Living Functional Materials. ACS Synth. Biol. 4, 8–11 (2015).

3. Walker, K. T., Goosens, V. J., Das, A., Graham, A. E. & Ellis, T. Engineered cell-to-cell signalling within growing bacterial cellulose pellicles. Microb. Biotechnol. 12, 611–619 (2019).

4. Limoli, D. H., Jones, C. J. & Wozniak, D. J. Bacterial Extracellular Polysaccharides in Biofilm Formation and Function. Microbiol. Spectr. 3, (2015).

5. Augimeri, R. V., Varley, A. J. & Strap, J. L. Establishing a Role for Bacterial Cellulose in Environmental Interactions: Lessons Learned from Diverse Biofilm-Producing Proteobacteria. Front. Microbiol. 6, 1282 (2015).

6. Reiniati, I., Hrymak, A. N. & Margaritis, A. Recent developments in the production and applications of bacterial cellulose fibers and nanocrystals. Crit. Rev. Biotechnol. 37, 510–524 (2017).

7. Fijałkowski, K., et al. Increased water content in bacterial cellulose synthesized under rotating magnetic fields. Electromagn. Biol. Med. 36, 192–201 (2017).

8. Choi, S. M. & Shin, E. J. The Nanofication and Functionalization of Bacterial Cellulose and Its Applications. Nanomaterials. 10, 406 (2020).

9. Gorgieva & Trček. Bacterial Cellulose: Production, Modification and Perspectives in Biomedical Applications. Nanomaterials. 9, 1352 (2019).

10. Gullo, M., La China, S., Falcone, P. M. & Giudici, P. Biotechnological production of cellulose by acetic acid bacteria: current state and perspectives. Appl. Microbiol. Biotechnol. 102, 6885–6898 (2018).

11. Klemm, D. et al. Nanocelluloses as Innovative Polymers in Research and Application. in Polysaccharides II (ed. Klemm, D.,) 49–96 (Springer, 2006). doi:10.1007/12_097.

12. Gilbert, C. et al. Living materials with programmable functionalities grown from engineered microbial co-cultures. bioRxiv 2019.12.20.882472 (2019) doi:10.1101/2019.12.20.882472.

13. Gilbert, C. & Ellis, T. Biological Engineered Living Materials: Growing Functional Materials with Genetically Programmable Properties. ACS Synth. Biol. 8, 1–15 (2019).

14. Savitskaya, I. S., Shokatayeva, D. H., Kistaubayeva, A. S., Ignatova, L. V. & Digel, I. E. Antimicrobial and wound healing properties of a bacterial cellulose based material containing *B. subtilis* cells. Heliyon 5, e02592 (2019).

15. Heinrich, M. K. et al. Constructing living buildings: a review of relevant technologies for a novel application of biohybrid robotics. J. R. Soc. Interface 16, 20190238 (2019).

16. Lombardo, D., Calandra, P., Pasqua, L. & Magazù, S. Self-assembly of Organic Nanomaterials and Biomaterials: The Bottom-Up Approach for Functional Nanostructures Formation and Advanced Applications. Mater. Basel Switz. 13, (2020).

17. Laromaine, A., et al. Free-standing three-dimensional hollow bacterial cellulose structures with controlled geometry *via* patterned superhydrophobic–hydrophilic surfaces. Soft Matter 14, 3955–3962 (2018).

18. Greca, L. G., Lehtonen, J., Tardy, B. L., Guo, J. & Rojas, O. J. Biofabrication of multifunctional nanocellulosic 3D structures: a facile and customizable route. Mater. Horiz. 5, 408–415 (2018).

19. Schaffner, M., Rühs, P. A., Coulter, F., Kilcher, S. & Studart, A. R. 3D printing of bacteria into functional complex materials. Sci. Adv. 3, eaao6804 (2017).

20. Hu, Y. & Catchmark, J. M. Formation and characterization of spherelike bacterial cellulose particles produced by *Acetobacter xylinum* JCM 9730 strain. Biomacromolecules 11, 1727–1734 (2010).

21. Florea, M. et al. Engineering control of bacterial cellulose production using a genetic toolkit and a new cellulose-producing strain. Proc. Natl. Acad. Sci. 113, E3431–E3440 (2016).

22. Gullo, M., La China, S., Petroni, G., Di Gregorio, S. & Giudici, P. Exploring K2G30 Genome: A High Bacterial Cellulose Producing Strain in Glucose and Mannitol Based Media. Front. Microbiol. 10, (2019).

23. Singhsa, P., Narain, R. & Manuspiya, H. Physical structure variations of bacterial cellulose produced by different *Komagataeibacter xylinus* strains and carbon sources in static and agitated conditions. Cellulose. 25, 1571–1581 (2018).

24. Czaja, W., Romanovicz, D. & Brown, R. malcolm. Structural investigations of microbial cellulose produced in stationary and agitated culture. Cellulose 11, 403–411 (2004).

25. Ryngajłło, M., Jacek, P., Cielecka, I., Kalinowska, H. & Bielecki, S. Effect of ethanol supplementation on the transcriptional landscape of bionanocellulose producer *Komagataeibacter xylinus* E25. Appl. Microbiol. Biotechnol. 103, 6673–6688 (2019).

26. Hu, Y., Catchmark, J. M. & Vogler, E. A. Factors impacting the formation of sphere-like bacterial cellulose particles and their biocompatibility for human osteoblast growth. Biomacromolecules. 14, 3444–3452 (2013).

27. Ford, T. J. & Silver, P. A. Synthetic biology expands chemical control of microorganisms. Curr. Opin. Chem. Biol. 28, 20–28 (2015).

28. Manzoni, R., Urrios, A., Velazquez-Garcia, S., de Nadal, E., & Posas, F. Synthetic biology: insights into biological computation. Integr. Biol. Quant. Biosci. Nano Macro. 8, 518–532 (2016).

29. Boo, A., Ellis, T., & Stan, G.-B. Host-aware synthetic biology. Curr. Opin. Syst. Biol. 14, 66–72 (2019).

