## Supplementary Material for "Bacterial cellulose spheroids as building blocks for 2D and 3D engineered living materials"

**Caro-Astorga *et al.* 2020**

**Supplementary Table 1.** Culture condition variables tested to find the determinant parameters for BC spheroids production. Not all combinations of parameters were tested. Only the container and the $OD_{600}$ of inoculation bacteria at the start of the culturing had a critical effect.

| Variable | Parameters |
|---|---|
| **Inoculum age** | 1-10 days |
| **Bacterial sub-culturing** | 1-4 subcultures |
| **Container** | 14, 15, 50 ml plastic tubes; glass flasks of 100, 250, 500 and 1000 ml with plain and irregular wall; glass bottles of 100, 250, 500 and 1000 ml |
| **Temperature** | 30°C and 25°C |
| **Medium:air ratio in container** | 1:3, 1:4, 1:5, 1:10 |
| **Shaking conditions** | 150, 250 and 350 rpm in incubator; 100 rpm with magnet stirrer |
| **Culture medium** | HS, 2x HS, +1% ethanol, + 8% glycerol, 1% fructose |
| **Time between seeding and shaking** | 0, 30 min, 1 hour and 2 hours |
| **Inclination of tubes** | 0, 15, 45, 65 and 90 degrees |
| **Preculture dilution** | 1:10, 1:100, 1:1000 |
| **Optical density of the final culture** | 0.1, 0.01, 0.001, 0.0001 |

**Supplementary video 1.** Time-lapse of *K. rhaeticus* cells growth for 2h in microfluidic plate with HS media supplemented with Fluorescence Brighter 28 to stain cellulose bands (blue). Bands of bacterial cellulose are produced continuously while bacteria continue the cell division process, resulting in branched cellulose bands. In the top area of the plate, a globular tangle of bacteria producing bacterial cellulose bands is forming the first stages of a pro-spheroid.

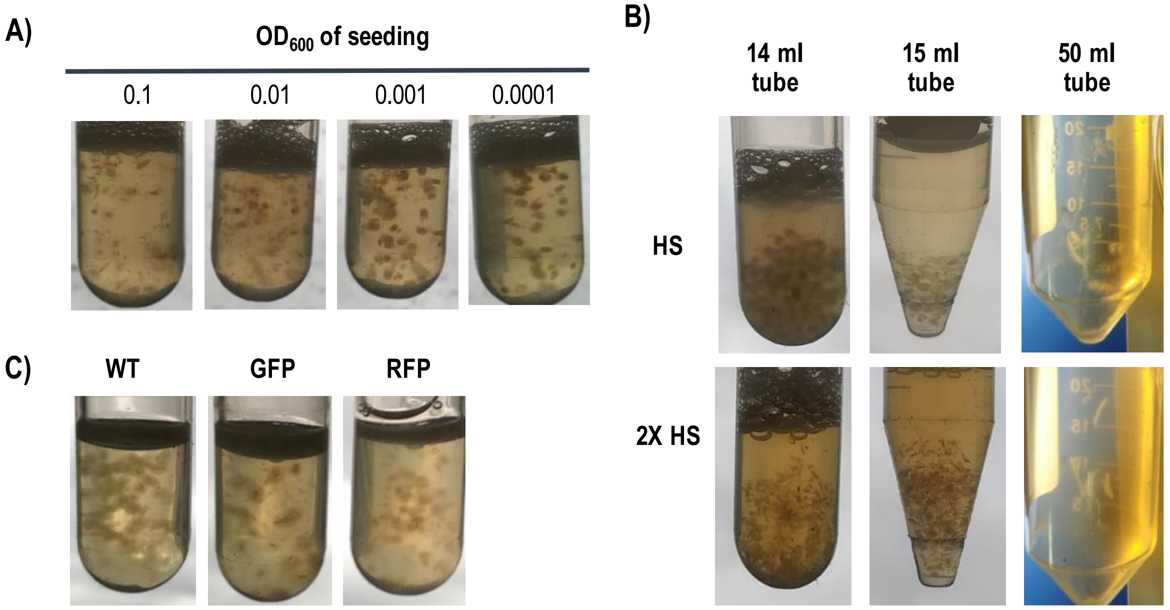


**Supplementary Figure 1.** BC spheroids growth conditions. A) Effect of Optical Density of the culture. Only OD below 0.001 produce a high yield spheroid. Higher densities produce fibrous material and interconnected spheroids. B) Effect of the shape of the tube and media concentration on spheroids production. 2x HS medium produce higher quantities of spheroids, but they were not of desired size. The shape of 50 ml tubes prevented spheroid production, with 14 ml culture tubes being the preferred choice. C) Cells of *K. rhaeticus* transformed with plasmid for high expression of sfGFP or RFP fluorescent proteins spheres produced similar spheroids than the wild type strain.

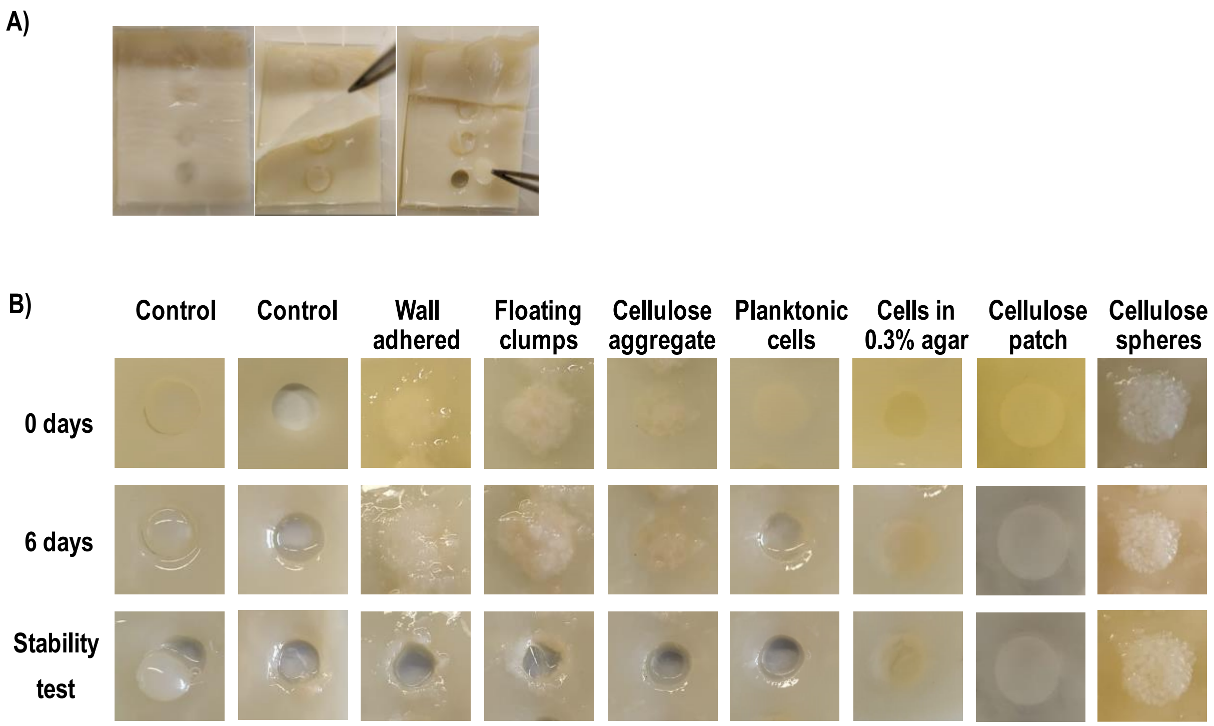


**Supplementary Figure 2.** A) Repair assay of a day 10 bacterial cellulose pellicle damaged with a hole puncture then incubated with static aeration for 7 days at 30°C in fresh HS media supplemented with 2% glucose. Over the original BC, a new pellicle grows that is poorly attached to the original pellicle and can be removed easily with a tweezer. B) Repair assay of a day 7 bacterial cellulose pellicle damaged with a hole puncture. Images shown are representative of three replicates. Upper line shows images of seeding the repairing bacteria. Middle and bottom line shows images after 6 days of incubation and after the stability test, respectively. Cells in different physiological state were used to heal the damage and incubated for 6 days at 30°C. The source of cells were: i) fragments of biofilm found adhered to the wall of a flask after 4 days in shaking conditions; ii) floating clumps formed in shaking conditions from a initial culture set with high cell density ($OD_{600}$ ~0.5); iii) cellulose aggregates present in the culture medium under the pellicle; iv) a pellet of cells grown in shaking conditions with cellulase, centrifuged, washed with HS and centrifuged again; v) cells from (iv) embedded in a 0.3% HS agar matrix at 40°C and immediately placed in the pellicle before it solidifies; vi) a cellulose patch of slightly bigger dimensions than the hole produced in the pellicle, used to force the edges of the patch and the hole to be in close contact; vii) cellulose spheroids from a day 3 shaking culture.
